# The dengue virus non-structural protein 1 (NS1) use the scavenger receptor B1 as cell receptor in human hepatic and mosquito cells

**DOI:** 10.1101/871988

**Authors:** Ana C. Alcalá, José L. Maravillas, David Meza, Octavio T. Ramirez, Juan E. Ludert, Laura A. Palomares

## Abstract

Dengue is the most common virus disease transmitted to humans by mosquitoes. The dengue virus NS1 is a multifunctional protein that form part of replication complexes. In addition, NS1 is also secreted, as a hexamer, to the extracellular milieu. Circulating NS1 has been associated with dengue pathogenesis by several different mechanisms. Cell binding and internalization of soluble NS1 result in the disruption of tight junctions and in down regulation of the innate immune response. In this work, we report that the HDL scavenger receptor B1 (SRB1) in human hepatic cells, and a scavenger receptor B1-like in mosquito C6/36 cells act as cell surface binding receptor for dengue virus NS1. The presence of the SRB1 on the plasma membrane of C6/36 cells, as well as in Huh-7 cells, was demonstrated by confocal microcopy. Internalization of NS1 can be efficiently blocked by anti-SRB1 antibodies and previous incubation of the cells with HDL significantly reduces NS1 internalization. In addition, the transient expression of SRB1 in Vero cells, which lack the receptor, renders these cells susceptible to NS1 entry. Direct interaction between soluble NS1 and the SRB1 in Huh7 and C6/36 cells was demonstrated in vivo by proximity ligation assays an in vitro by surface plasmon resonance. Finally, data is presented indicating that the SRB1 also act as cell receptor for zika virus NS1. These results demonstrate that dengue virus NS1, a *bona fide* lipoprotein, usurps the HDL receptor for cell entry and offers explanations for the altered serum lipoprotein homeostasis observed in dengue patients.

## Introduction

The *Flavivirus* genus, belonging to the *Flaviviridae* family, includes a group of vector borne viruses of importance in human public health as dengue virus (DENV), Zika virus (ZIKV), yellow fever virus (YFV), Japanese encephalitis virus (JEV) and West Nile virus (WNV). DENV and ZIKV are transmitted by the same arthropods of the *Aedes* genus and share a similar geographical distribution (Paixão et al 2018). It is estimated that almost half of the world population, distributed in tropical and subtropical zones, is at risk of contracting these diseases (WHO, 2018). The initial symptoms of both DENV and ZIKV infections manifest as a febrile syndrome. Infections with DENV can evolve to hemorrhagic syndrome accompanied of hypovolemic shock, systemic failure and death, mainly in children. In turn, ZIKV infections when acquired during pregnancy, are associated with the occurrence of microcephaly and other neurological severe injuries in the fetus brain (Rather et al 2017). In adults, ZIKV has been associated with the occurrence of Guillan-Barré syndrome which is highly disabling (Song et al., 2017). DENV and ZIKV virions are spherical particles of about 50nm in diameter. The positive RNA genome is about 11Kb in length encoding for 3 structural (Membrane, Capsid and Envelope) and for 7 nonstructural proteins (NS1, NS2A, NS2B, NS3, NS4A, NS4B and NS5). The gene encoding for the NS1 is 1056 ntds in length rendering a 48-54 KDa protein depending on its glycosylation status. The NS1 could be found in monomeric, dimeric or hexameric forms (Muller and Young 2013). Structurally, the monomeric form of NS1 is composed of three domains: α/β wing (RIG-I-like folded, from amino acids 38–151), β-roll (hydrophobic, from amino acids 1–29) and β-ladder (composed of a β-sheet and a spaghetti loop, from aminoacids 181–352) (Akey et al., 2015). Once the NS1 is translated from the viral genome, it is located at the lumen of the ER in a monomeric form which rapidly dimerizes (reviewed in Muller and Young, 2013). The intracellular NS1 is mainly dimeric and is associated to intra and extracellular membranes participating in different processes interacting with cellular and viral proteins during the viral replication, virion assembly, signaling transduction and evasion of the cellular immune response (Jacobs et al., 2000; Scaturro et al., 2015). The hexameric form of NS1 is an association of three dimers assembled in a hollow barrel-shaped structure, with a central channel rich in lipids acquired from cellular membranes (Gutshe et al., 2011). NS1 is the only flaviviral nonstructural protein secreted together with the viral particles toward the extracellular milieu from infected vertebrate and insect cells in high concentrations (up to μg/mL) (Flamand et al., 1999; Alcala et al., 2016). In fact, NS1 is used as an early marker for DENV and ZIKV detection from patient serum samples (reviewed in De la Cruz Hernández et al., 2013). The DENV, soluble circulating NS1 has been involved in several pathogenic mechanisms during the infection process. It has been reported NS1 can affect the complement pathway promoting the degradation of C4 protein, alters the coagulation system inhibiting the activation of prothrombin (reviewed by Conde et al., 2017). NS1 can also disrupts endothelium integrity, promoting an increase in the vascular permeability after internalization by endocytosis, that may be related to plasma leakage (Glasner et al., 2017; Puerta-Guardo et al., 2019; Wang et al., 2019). Moreover, pre-exposure of human liver or dendritic cells to NS1 renders these cells more susceptible to DENV replication (Alcon-LePoder et al, 2005; Alayli and Scholle, 2016). Thus, a better understanding of the DENV soluble NS1- cell interactions may result in strategies to combat DENV replication and pathogenesis.

The scavenger receptor B type 1 (SRB1) or his human homolog CLA-1, is a 509 amino acid transmembrane glycoprotein containing an extracellular domain as a unique loop of 408 amino acids with multiple sites of N-glycosylation and a cysteine-rich region, 2 transmembrane domains of 22 and 23 amino acids and 2 cytoplasmatic N and C terminal domains of 9 and 44 amino acids, respectively (Shen et al, 2018). SRB1 is the physiologically relevant cell surface HDL receptor responsible for selective HDL-cholesteryl esters (HDL-CE) uptake mainly in liver, where it is most abundantly expressed, providing cholesterol for bile acid synthesis and controlling de plasma HDL levels (Acton et al., 1996; Shen, 2019). In insects as *Drosophila melanogaster* and *Aedes aegypti*, among others, homolog genes for SRB1 have been identified; however, their functions in fatty acid transport has not been studied in these organisms (Nichols et al., 2008).

The SRB1 has been described as the principal receptor of hepatitis C virus (HCV) in hepatocytes, through the recognition of the E2 hypervariable region 1 of the HCV envelope lipoprotein (Catanese et al., 2010). In addition, it has been reported that apolipoprotein AI (ApoA-I) the principal carrier of HDL molecules, bridges DENV particles and cell receptor SRB1 and facilitates entry of DENV into cells enhancing the infection (Li et al., 2013). DENV has a broad tropism for liver tissue where it replicates efficiently in hepatocytes; however, the elements that direct this tropism are not known (Povoa et al., 2014). DENV NS1 can entry into liver tissue cells both *in vitro* and *in vivo* in the absence of viral particles. In addition, an increase in virus yield is observed from hepatic cells preincubated with DENV NS1, and it has been postulated that DENV NS1 conditions the hepatic cells to increase viral particle production (Alcon-Le Poder et al., 2005). Recently, it was reported that endocytosis of NS1 is required for NS1-mediated endothelial hyperpermeability (Wang et al., 2019). However, the host factors favoring NS1 binding and entry in hepatocytes or other cells are unknown.

Since NS1 must be internalized to exert its functions and the DENV NS1 is a *bonna-fide* glyco-lipoprotein, with a lipid cargo composed of triglycerides, cholesteryl esters and phospholipids that resembles in composition the plasma lipoprotein HDL (Gutshe et al., 2011), this suggested to us that DENV NS1 could mimic or hijack some of the lipid metabolic pathways to enter the cell. In this work, we present evidence demonstrating that the HDL SRB1 in human hepatic cells, and an SRB1-like molecule in mosquito cells, is also used by DENV and ZIKV soluble NS1 as cell receptor for cell binding and internalization.

## Material and Methods

### Virus and cells

Dengue 4 virus (kindly donated by Dr. Juan Salas, Instituto Politécnico Nacional, México) was propagated in suckling mouse brains as previously described (Gould and Cleg, 1991). Zika virus strain M7366 (kindly donated by Dr. Susana López, Instituto de Biotecnología, UNAM, Mexico) was propagated in C6/36 cells. Virus titers from mice brain homogenates or cell culture supernatant were determined by plaque assay as described by Ludert et al. (2008) but using Vero cells grown in the cells culture conditions described in this work. C6/36 cells (ATCC, USA) were grown in EMEM medium at 28 °C and 5% CO_2_ atmosphere. Huh-7 and Vero cells were grown in DMEM medium (Sigma, USA) at 37 °C and 8% CO_2_ atmosphere. All cell culture media were supplemented with 5% FBS.

### Cell surface localization of SBR1

Confluent monolayers grown in LAB-TEK^®^ (Nalgene Nunc International, USA) were fixed (4% paraformaldehyde-PBS) for 10 min at RT and then incubated with blocking buffer (10% FBS, 3%BSA, 100mM Glycine in PBS) for 45 min at RT. The primary antibodies anti-SRB1 rabbit Mab (Abcam, USA) was diluted 1:300 (10% FBS, 1% BSA, 100mM Glycine in PBS) and incubated with the cells for 1h, 37°C. After 5 washes, cells were incubated with an anti-rabbit Alexa Fluor 594 (Invitrogen, USA) diluted 1:1000 in PBS. After 10 washes, the cells were incubated with Dapi (4′,6-Diamidine-2′-phenylindole dihydrochloride) in mounting medium (Molecular Probes). Afterward, a coverslip was added and sealed with nail polish. The immunolabeled cells were observed in the inverted confocal microscope (Olympus MPhot) and the images were analyzed in the Image J/FIJI software. In order to capture images with higher resolution of the molecules located on the plasma membranes, monolayers of Huh-7 and C6/36 were cells grown in Fluorodish Cell Cultures (WPI, USA) and the SRB-1 coimmunolabelled with wheat germ agglutinin conjugated with Alexa Fluor488 (Thermofisher, USA) to observed the silhouette of the cells and the integrity of the plasmatic membrane. Colabelled cells were examined by total internal reflection fluorescence microscopy (TIRF) (Olympus IX81 TIRF). The antibodies and conditions used in TIRF were as described for the confocal microscopy.

### NS1 internalization assays

Huh-7 and C6/36 cells were grown in 8 well chambers (LAB-TEK^®^, Nalgene Nunc International, USA) for 24h, until confluent monolayers were formed. Monolayers were incubated with 3,5 μg/well of recombinant DENV or ZIKV NS1 protein (Aalto Bio Reagents, Dublin). After 1,5h incubation, at the cell specific growth temperature, cells were washed extensively, fixed with 4% paraformaldehyde-PBS for 10 min at RT and permeabilized with 0,1 % Triton-PBS, for 5 min at RT and finally incubated with blocking buffer (10% FBS, 3% BSA, 100mM Glycine in PBS) for 45 min at RT. Internalized NS1 was revealed by immunolabeling using an anti-NS1 Mab (kindly donated by Prof. Eva Harris, University of California, Berkeley) diluted 1:300 (10% FBS, 1% BSA, 100mM Glycine in PBS) as primary antibody incubated for 1h at 37°C. After 5 washes, cells were incubated with an anti-mouse Alexa fluor 568 (Invitrogen, USA) diluted 1:1000 in PBS as secondary antibody. After 10 washes, the cells were incubated with Dapi (4′,6-Diamidine-2′-phenylindole dihydrochloride) in mounting medium (Molecular Probes). Afterward, a coverslip was added and sealed with nail polish. The immunolabeled cells were observed in an inverted confocal microscope (Olympus MPhot) and images analyzed with Image J/FIJI software. For the quantification of NS1 levels inside the cells, 10 cells by field on 3 fields by condition, were selected as Region of Interest (ROI), and arbitrary fluorescent units were measured using the Image J/Fiji software. The means for each condition were calculated and the analysis of variance (ANOVA) one-tailed test used to compare the mean of fluorescent arbitrary units (FAU) from each measure.

### DENV and ZIKV infection in insect cells preincubated with recombinant NS1

C6/36 cell monolayers grown in 24-well plates were preincubated or not with 3,5 and 7,0 μg/well of recombinant DENV or ZIKV NS1 protein for 1,5 h. After extensive washing, cells were inoculated with DENV or ZIKV at a MOI=1. Infection was left to proceed for 48 hours, the supernatants collected, and the virus yield titrated by plaque assay using Vero cells, as previously described (Ludert et al., 2008).

### Inhibition of NS1 entry by preincubation with anti-SRB1 antibodies

Confluent monolayers of Huh-7 and C6/36 cells, grown in 8-well chambers LAB-TEK^®^ (Nalgene Nunc International, USA) for 24h, were preincubated or not with anti-SRB1 antibody (Abcam) at concentrations of 1 and 10 μg/ml for 1h at 37°C. After extensive washing, cells were incubated with recombinant DENV NS1 protein at a concentration of 3,5 μg/well for 1,5 h, and the cells fixed and processed for detection and quantification of internalized NS1 as described above.

### NS1 and HDL competition assays

C6/36 or Huh7 were seeded in 8-well chambers and confluent monolayers were incubated with 150 or 300 μg/mL of HDL (Merck, USA) in specific growth medium without FBS for 1.5 h, at the appropriate cell growth conditions. After extensive washing, 3,5 μg/well of DENV or ZIKV recombinant NS1 protein were added in growth medium without FBS. After incubation for 1,5h at the cell specific growth conditions, cells were fixed, permeabilized and immunolabeled for detection of internalized DENV or ZIKV NS1 as describe above.

### Overexpression of SRB1 in Vero cells

Monolayers, 80-90% of confluence, of Vero cells seeded in 8-well chambers, were transfected with the plasmid pCMV-SCARB1-His (Sinobiological, USA) for the expression of the SRB1. Transfections were carried with Lipofectamine 2000 LTX (Invitrogen) according to manufacturer’s instructions using 0.5 μg/well of the plasmid. Twenty-four hpt, supernatant were removed and the cells incubated with 3,5 μg/mL of recombinant DENV NS1 in DMEM medium without FBS for 1.5h at the cells growth conditions. Cells were fixed, permeabilized and co-immunolabeled with anti-NS1 and anti-SRB1 specific primary antibodies and reveled and visualized by confocal microscopy as describe above.

### Proximity Ligation Assays

**I**nteractions between the SRB1 and NS1 proteins from DENV and ZIKV in Huh-7 and C6/36 cells were detected using the commercial kit Duolink (Sigma-Aldrich) based on *in situ* proximity ligation assay (PLA) used following manufacturer’s instructions. Briefly, C6/36 or Huh-7 monolayers were grown on 8-well chambers, preincubated with DENV or ZIKV NS1 recombinant proteins (3,5 μg/well), and after 45 min of incubation, fixed with 4 % paraformaldehyde for 10 min at RT. After incubation with blocking buffer for 30 min at 37 °C, cells were incubated with mouse anti-NS1 mAb and rabbit anti-SRB1 polyclonal antibody (Abcam, USA) as primary antibodies. After several washes, cells were incubated with the PLA PROBES linked, anti-mouse and anti-rabbit secondary antibodies. Finally, the cells were subjected to ligation and amplification reactions. The specificity of PLA was evaluated using recombinant NS1 exposed cells incubated only with the NS1 specific antibody and the two PLA PROBES, as negative technical control. The PLA signal was detected in a confocal microscope (Inverted Olympus MPhot) using a 60X immersion oil objective and values determined considering the mean of three images per condition in three independent experiments. Image analysis was carried out using the Fiji ImageJ software, and the signal quantified as described in Alcalá et al. (2017).

### Surface Plasmon Resonance

Recombinant, human SRB1 (Sino Biological, USA) was covalently immobilized onto carboximethylated-dextran sensor chips (CM5 series S, General Electric Healthcare). Previously, the sensor chip surface was activated using 1-ethyl-3-(3 dimethylaminopropyl) carbodiimide (EDC) and N-hydroxysuccinimide (NHS) reagents, according to the standard procedure for amine coupling. For direct capture-coupling to NH_2_ (in immobilization buffer of 10 mM sodium acetate, pH 5.0) the SRB1 was diluted to 30 μg/mL and injected at 5 μg/min flow-rate until reach a density of 10,000 resonance units (RU), using running buffer HBS-EP*. The remaining surface-active groups were blocked with 1M ethanolamine hydrochloride, pH 8.5. A dilution series of the DENV recombinant NS1 protein (Aalto Bio Reagents, Dublin), at six concentrations of 23.4, 46.8, 93.7, 187.5, 375 y 750 nM, were then injected at a flow rate of 30 μL/min for 200s of association with SR-B1, followed by dissociation step adjusted to 600s. The surface was successively regenerated between analyte injections with a single-brief (30s) injection of glycine-HCl, pH 2.5. All analyses were carried out at 25° C using a Biacore T200 (General Electric Healthcare). Surface plasmon resonance (SPR) response data defined as sensorgrams, were zeroed at the beginning of each individual injection and then double referenced. The responses were plotted against the analyte concentration and fit to a 1:1 (A+B = AB) binding model using Biacore T200 evaluation software, version 2.0 (General Electric Healthcare).

## Results

### A scavenger-like receptor is expressed insect cells plasma membrane

The expression of SRB1 in liver cells has been widely reported (Shen et al 2019). To evaluate the expression of SRB1 in C6/36 cells, non-permeabilized C6/36 monolayers were immunolabeled with an antibody specific for human SRB1 and analyzed by confocal microscopy, using Huh-7 and Vero cells as controls. The results indicated that the SRB1 is expressed in the cell membrane of Huh-7 and C6/36 cells, but not in the kidney derived Vero cells (Fig. 1A). To further corroborate that C6/36 cells indeed express the SRB1 on the cell surface, the location of the SRB1 on C6/36 was determined by TIRF microscopy, using Huh-7 as positive controls. As shown in Figure 1B, a clear signal for the SRB1 was obtained, indicating that C6/36 cells express a SRB1 molecule on the plasma membrane. Finally, it was inquired if a sequence for this receptor was annotated in the genome of *Aedes albopictus*; a putative gene for SRB1, encoding a protein with and estimated molecular weight of 58 Kd, and a homology of 33.4 % with the human protein, was found in the NCBI protein data base, under the code XP_019526049.2. These results taken together indicate that C6/36 cells express an SRB1-like protein on the plasma membrane.

**Figure 1.**
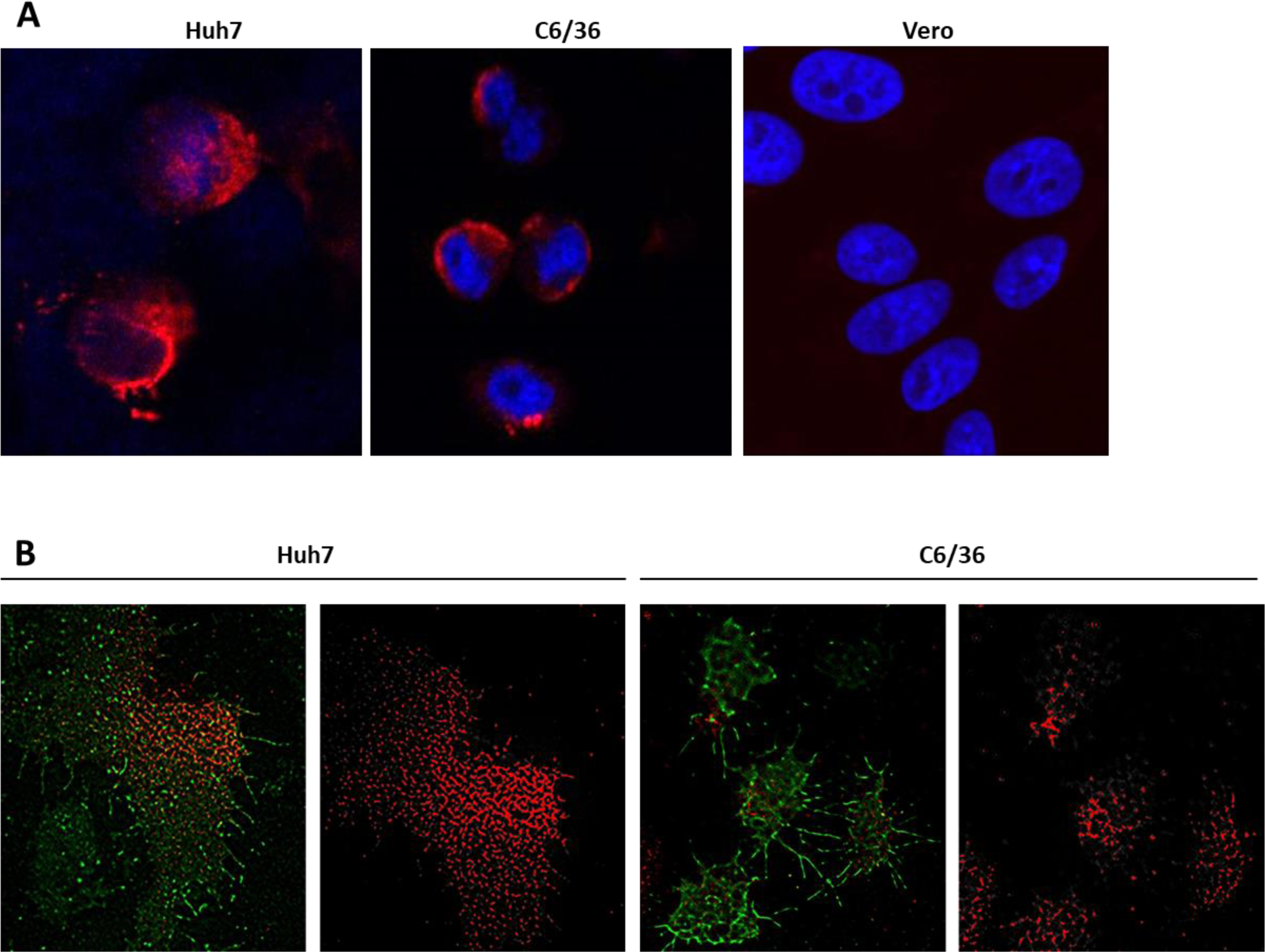
Detection of the scavenger receptor B1 (SRB1) in liver, mosquito and kidney derived cell lines. A) Confluent cell monolayers were fixed, stained for SRB1 (red), the nuclei counterstained with DAPI (blue) and observed by confocal microscopy. B) The presence of the SRB1 on the cell plasma membrane was corroborated by Total Internal Reflection Fluorescence (TIRF) Microscopy, costainning cells for SRB1 (red) and wheat germ agglutining (green) as cell membrane marker.

### DENV and ZIKV NS1 are efficiently internalized in hepatic and insect cells

It is well established that DENV NS1 is internalized in human hepatic and dendritic cells, rendering the cells more susceptible to DENV infection (Alcon-LePoder et al, 2005; Alayli and Scholle, 2016); yet internalization of DENV NS1 in mosquito cells or of ZIKV NS1 in any cell type have not been reported. Thus, using Huh-7 liver cells in parallel, we evaluated if DENV and ZIKV NS1 were also internalized by C6/36 mosquito cells after 1 or 2 hours of incubation. As shown in Figure 2, permeabilized cells were readily stained with anti-NS1 antibodies indicating that DENV and ZIKV NS1 were efficiently internalized by both cell lines. However, different fluoresce patters were observed for the internalized DENV NS1 in liver and mosquito cells; while a punctuate pattern was observed in liver cells, a diffuse cytoplasm distribution for NS1 was observed in the mosquito cells. Likewise, different fluoresce patters were observed between internalized DENV and ZIKV NS1 in liver cells, with ZIKV NS1 showing a fiber-like pattern. These observations, in agreement with previous data (Puerta-Guardo, et al., 2019), suggest that there may be cell and virus dependent differences in the way NS1 is internalized.

**Figure 2.**
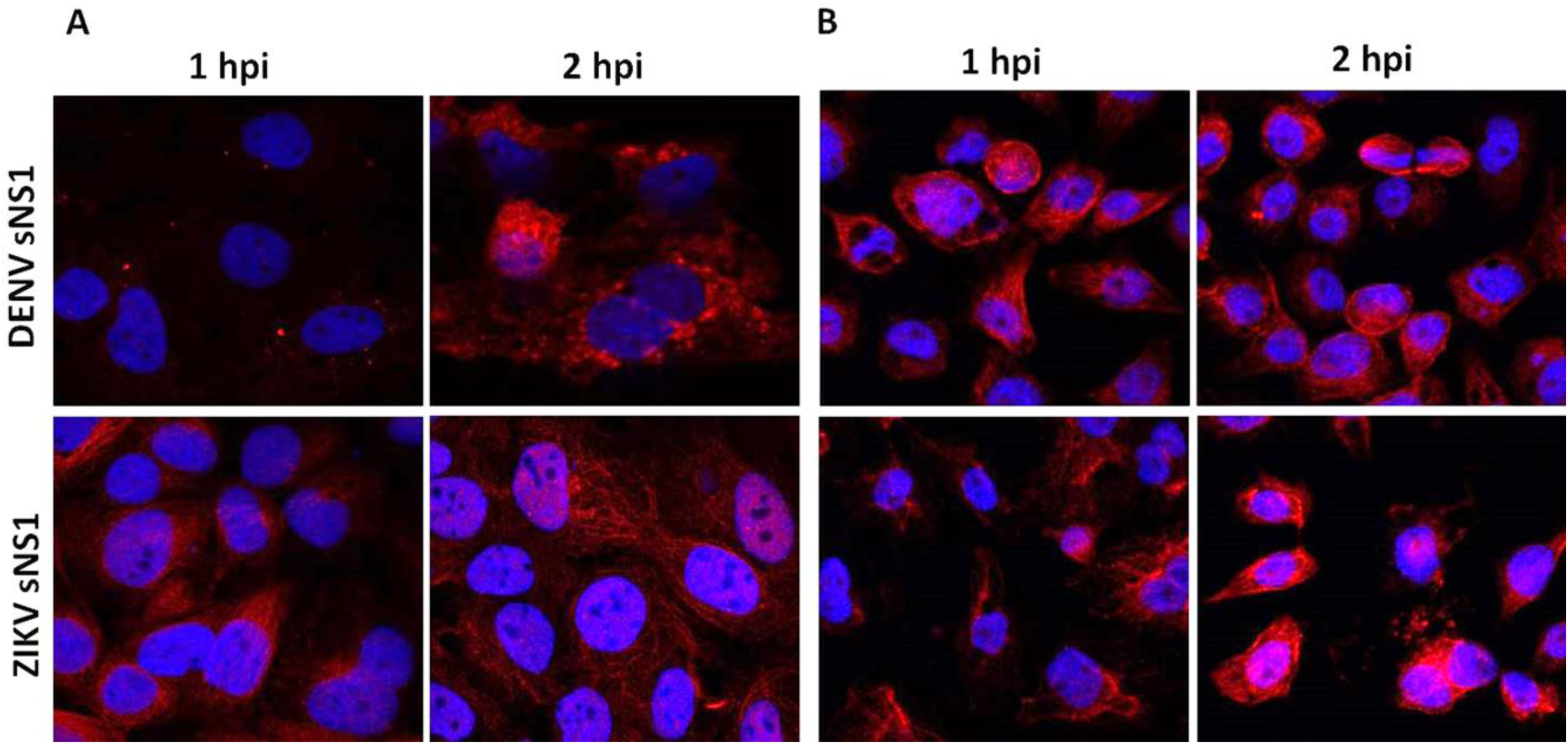
Internalization of DENV and ZIKV rNS1 in Huh-7 (A) and C6/36 cells (B). Cells were preincubated with recombinant NS1 and at the indicated times cells were washed, fixed, permeabilized and stained for intracellular NS1 (red), nuclei counterstained with DAPI and observed by confocal microscopy.

### DENV and ZIKV NS1 enhance the viral yield in insect cells

Once the entry of NS1 into C6/36 and Huh7 cells was assessed, next whether preincubation of mosquito cells with ZIKV or DENV NS1 will render the cells more susceptible to virus replication was evaluated. For this, C6/36 confluent monolayers were incubated with 0, 3.5 or 7 μg/well of NS1 for 1,5 h. Afterward, cells were infected with DENV or ZIKV at a MOI = 1 and at 48 hpi the supernatants were collected to determine the viral titer. A significant increase in the virus yield was observed from DENV and ZIKV NS1 pre-incubated cells in comparison with those not pre-incubated (Fig 3). The viral titers for DENV were 2.5×10^3^, 6.5×10^3^ and 7×10^3^ UFP/mL and for ZIKV, 4.5×10^4^, 1.5×10^5^ and 1.8×10^5^ UFP/mL in cells pre-exposed to 0, 3.5 and 7 μg/mL of NS1, respectively. The viral titer did not experience a significant increase in cells pre-incubated with 3.5 ug/mL or 7 ug/mL of NS1, perhaps due to a saturation of the system. This result suggests that exposure and internalization of NS1 by C6/36 cells before viral infection favors viral particles production, in agreement with previous results reported for liver and dendritic cells (Alcon-Le Poder et al., 2005; Alayli and Scholle, 2016).

**Figure 3.**
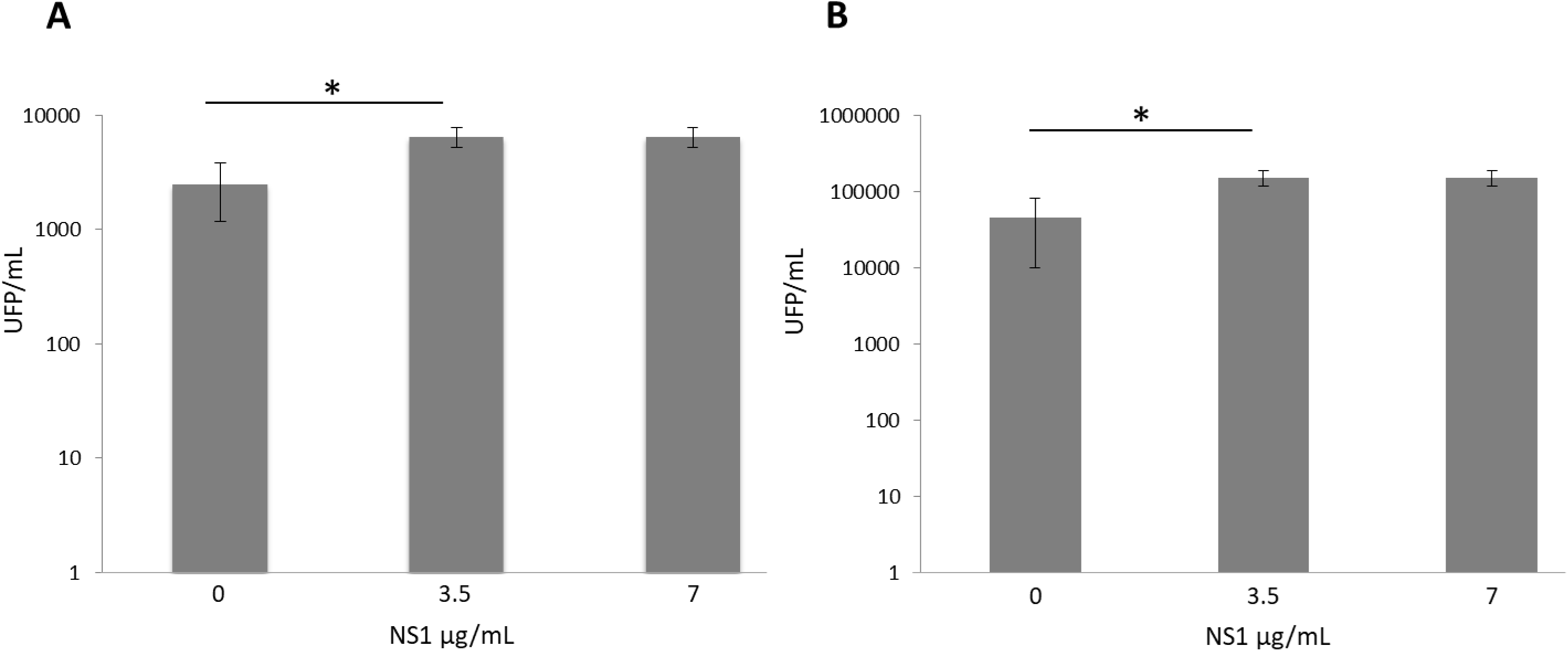
DENV (A) and ZIKV (B) titers after preincubation or not of C6/36 cells with recombinant NS1. Monolayers of C6/36 cell were preincubated for 1,5h with the indicated concentrations of either DENV or ZIKV NS1. After washing, cells were infected with the corresponding virus at a MOI=3 for 1h, infection allowed to proceed and the supernatants collected at 24 hpi, titrated by plaque assays in Vero cells. * *p*=0.05.

### SRB1 participate in the entry of NS1 in hepatic and insect cells

To evaluate the participation of the SRB1 in the entry of NS1, monolayers of C6/36 or Huh7 cells were treated with 0, 1 and 10μg/mL of anti-SRB1 monoclonal antibody to block the SRB1. After removal of the antibody, cells were incubated with DENV NS1. Cells were fixed, permeabilized and immunolabeled to detect internalized DENV NS1 as described above. Fluorescent signals were quantified to compare the internalization of the NS1 in the presence or absence of the antibody. A significant decrease of approximately 30%, in the intracellular specific signal for NS1 was observed in Huh7 cells pre-incubated with the specific monoclonal antibody compared to controls, no pre-incubated cells. No significant difference in the signal was observed when the cells were pre-incubated with 1 or 10 μg/mL of the antibody (Fig. 4A and B). A significant reduction in the internalization of DENV NS1 was also observed in C6/36 cell treated with the anti-SRB1 antibody (Fig. 4C and D). The specific intracellular fluorescent signal for DENV NS1 in C6/36 cells, showed a decrease of approximately 38% and 63% when preincubated with 1 or 10 μg/mL of the anti-SRB1 antibody, respectively, in comparison with the control. The significant decrease in the intracellular NS1 specific signal observed in the preincubated cells suggests that blocking SRB1 with specific antibodies affects the entry of NS1 in both cell lines.

**Figure 4.**
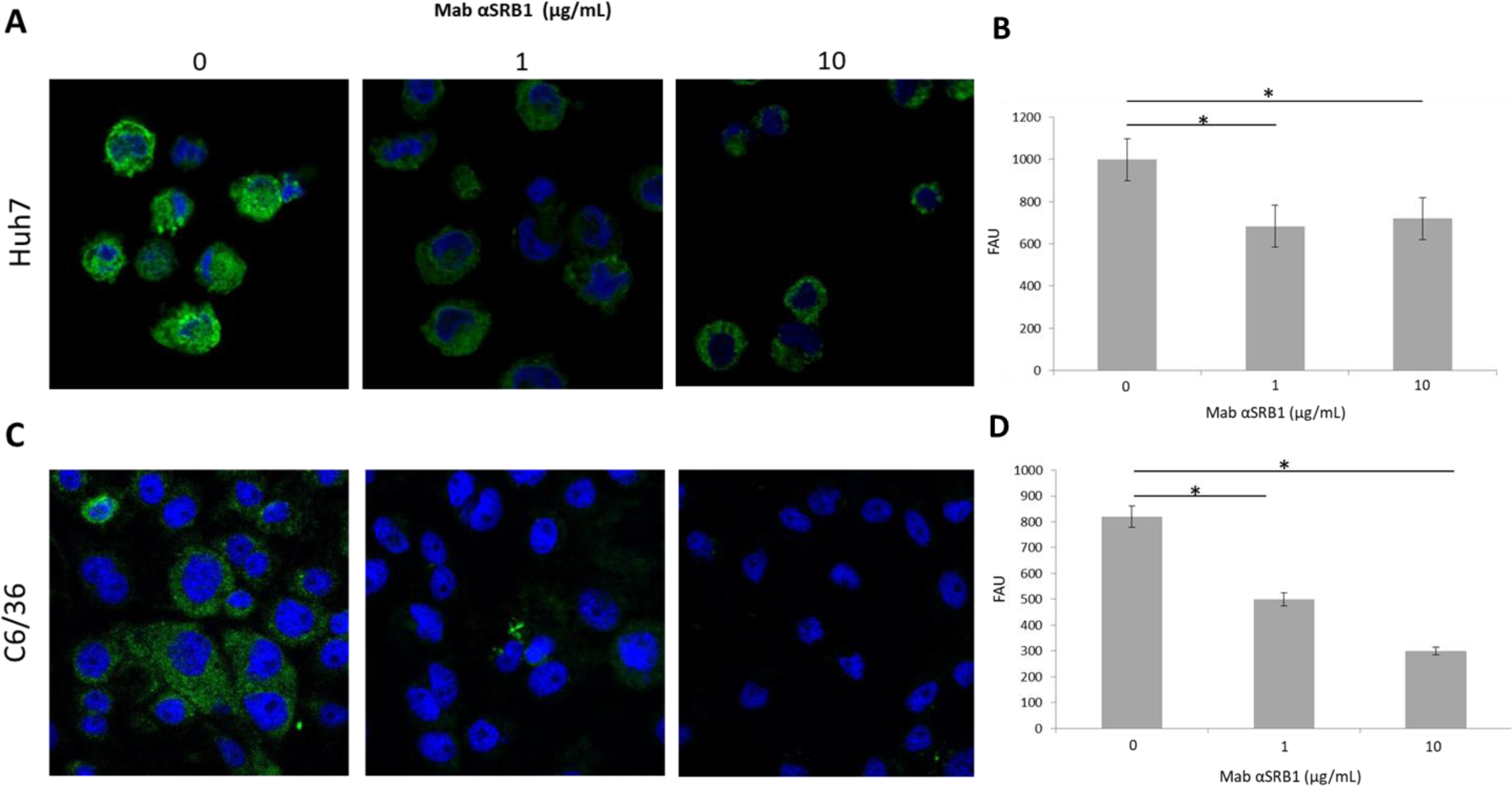
Inhibition of NS1 internalization by antibodies against the SRB1. Huh7 cells (A,B) and C6/36 (C,D) cells were incubated with anti-SRB1 antibodies at the indicated concentrations for 1h. After extensive washing, cells were incubated with DENV and ZINK NS1 (3,5 μg/well) for 90 min. Finally, cells were fixed, permeabilized and labelled for NS1. Cells were analyzed by confocal microscopy (left panels). The levels of NS1 inside the cells and expressed as fluoresce arbitrary units (FAU) (right panels). Differences in FAU were compared for significance using an ANOVA test. * *p*=0.05. n=3.

HDL is the natural ligand of SRB1 in human liver cells. In order to evaluate if the natural ligand of the SRB1 will compete with NS1, monolayers of Huh-7 and C6/36 cells were pre-incubated with human HDL at 0, 75 or 150 μg/mL. After several washes to remove unbound HDL, the monolayers were incubated with 3,5 μg/well of DENV NS1 and for this competition studies, ZIKV NS1 was also included, at the same concentration. Huh-7 cells showed a reduction of approximately 28 and 35% in the amount of internalized DENV NS1. In addition, the incubation of Huh-7 cells with HDL also reduced the amount of internalized ZIKV NS1 in approximately 22 and 45%, depending on the HDL concentration (Fig. 5A). Interestingly, incubation of C6/36 cells with human HDL also caused significant reductions in the amount of internalized DENV and ZIKV NS1. With DENV NS1 reductions of approximately 64 and 76% were observed and with ZIKV NS1 reductions of approximately 81% and 86% were observed, depending on the HDL concentration (Fig, 5B). These results are in agreement with the results obtained with the anti-SRB1 antibodies and indicate that SRB1 act as the cell receptor for both DENV and ZIKV NS1 in vertebrate and mosquito cells. Of note, attempts to knock down the expression of the SRB1 in Huh-7 cells using siRNAs were unsuccessful since in our hands, transfection of the siRNAs specific for the SRB1, resulted highly toxic for these cells.

**Figure 5.**
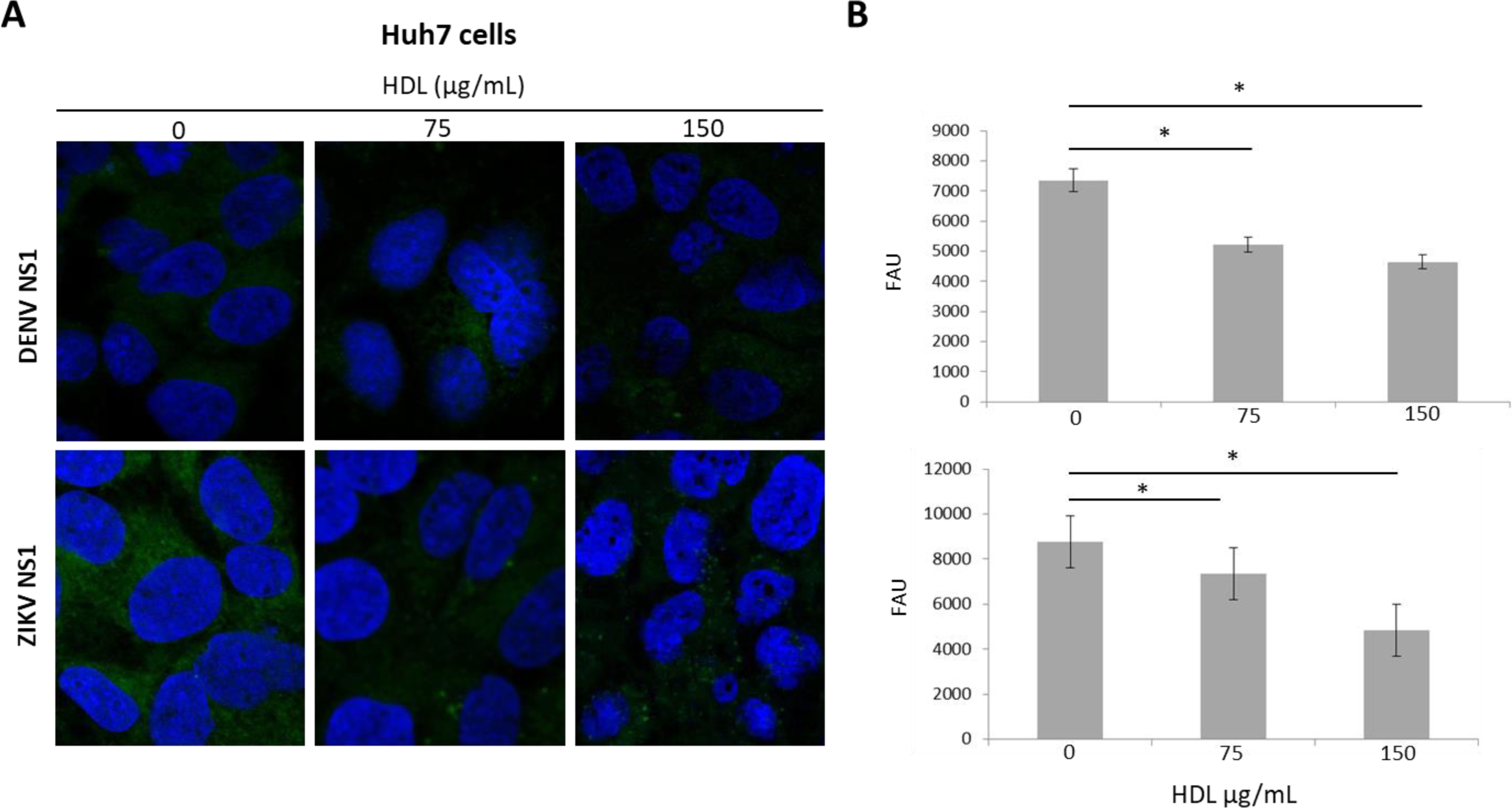
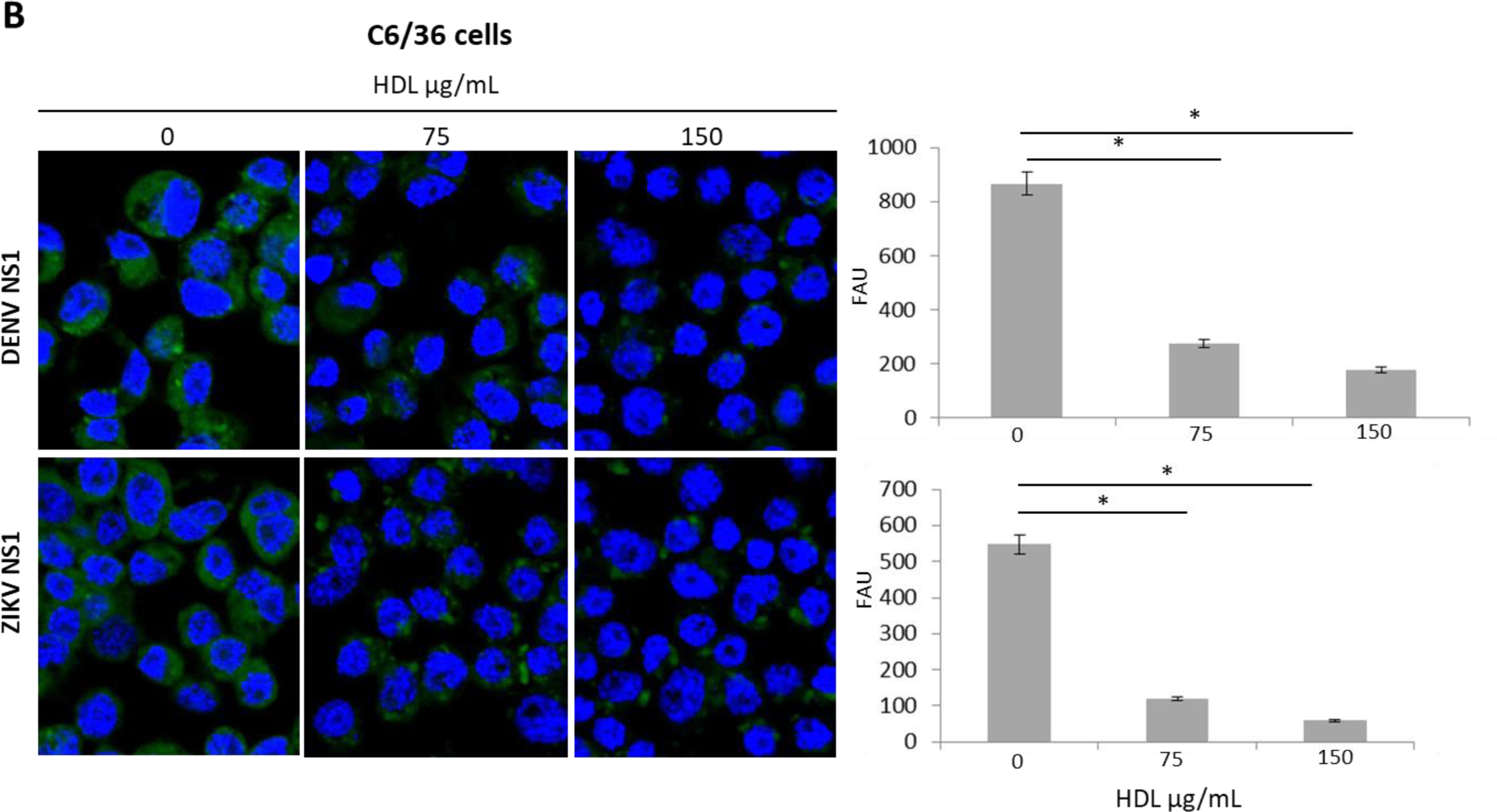
Competition assays between human HDL and recombinant DENV and ZIKV NS1. **A)** Huh-7 (A) and C6/36 cell (B) cells were incubated with HDL at the indicated concentrations for 90 min; after extensive washing, cells were incubated with DENV and ZINK NS1 (3,5 μg/well) for 90 min. Finally, cells were fixed, permeabilized and labelled for NS1. Cells were analyzed by confocal microscopy (left panels). The levels of NS1 inside the cells and expressed as fluoresce arbitrary units (FAU) (right panels). Differences in FAU were compared for significance using an ANOVA test. * *p*=0.05. n=3.

### Overexpression of the SRB1 in Vero Cells

We observed previously that Vero cells do not express the SRB1 (Fig. 1). In order to further demonstrate the participation of the SRB1 in the DENV NS1 internalization process, a gain of function experiment, overexpressing the SRB1 in Vero cells was conducted. Confluent monolayers of Vero cells were lipo-transfected with a plasmid expressing the SRB1. The expression of the SRB1 were determined at 24 hpt by confocal microscopy. Despite the different conditions tested to express the SRB1, the maximum percentage of cells expressing the SRB1 without cell damage, was around 7%, observed at 24 hpt when combining 0,5 μg/well of the plasmid with 2μL of transfectant reagent (Fig. 6). Under these conditions, cells were incubated with DENV NS1, fixed, permeabilized and simultaneously immunolabeled for SRB1 and DENV NS1. As shown in Figure 6, internalization of DENV NS1 was observed only on those cells expressing the SRB1. These data indicate that the sole expression of the SRB1 is enough to render Vero cells capable of internalizing DENV NS1.

**Figure 6.**
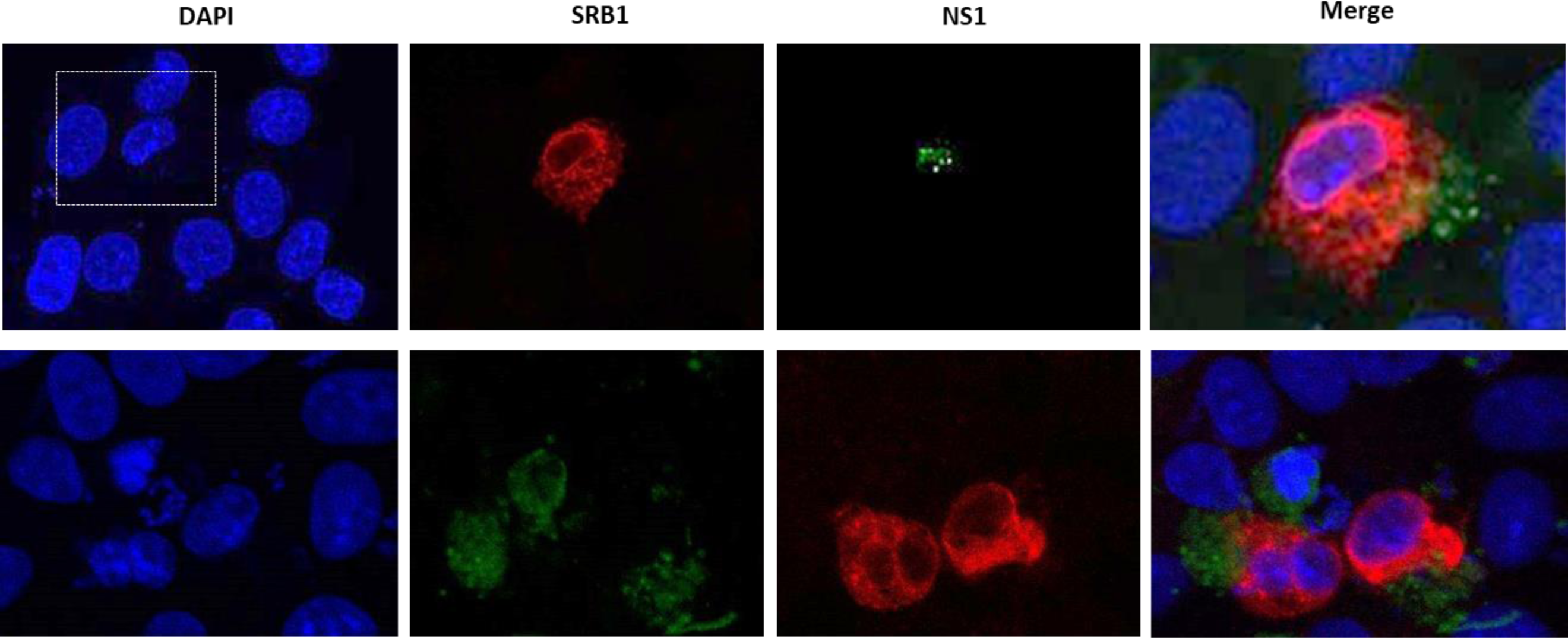
Expression of the SRB1 receptor in Vero cells. Vero cells were transfected with a plasmid encoding the SRB1 and 24 hpt incubated with recombinant DENV NS1 for 90 min. Cells were fixed, permeabilized and co-immunolabeled with anti-NS1 (green) and anti-SRB1 (red) and analyzed by confocal microscopy. Nuclei were counterstained with DAPI.

### SRB1 interacts with DENV and ZIKV NS1 protein *in vivo* and *in vitro*

To determine if there is actual physical interaction between DENV and ZIKV NS1 and the SRB1, the interactions between them on C6/36 and Huh7 cells was analyzed *in situ* by PLA. The assay clearly revealed that the DENV and ZIKV NS1 interact with the SRB1 in both cell lines (Fig. 7A, C, E and G). Meanwhile, no signal was observed in the technical controls run in parallel (Fig. 7 B, D, F, H), showing the specificity of the red dots signal observed.

**Figure 7.**
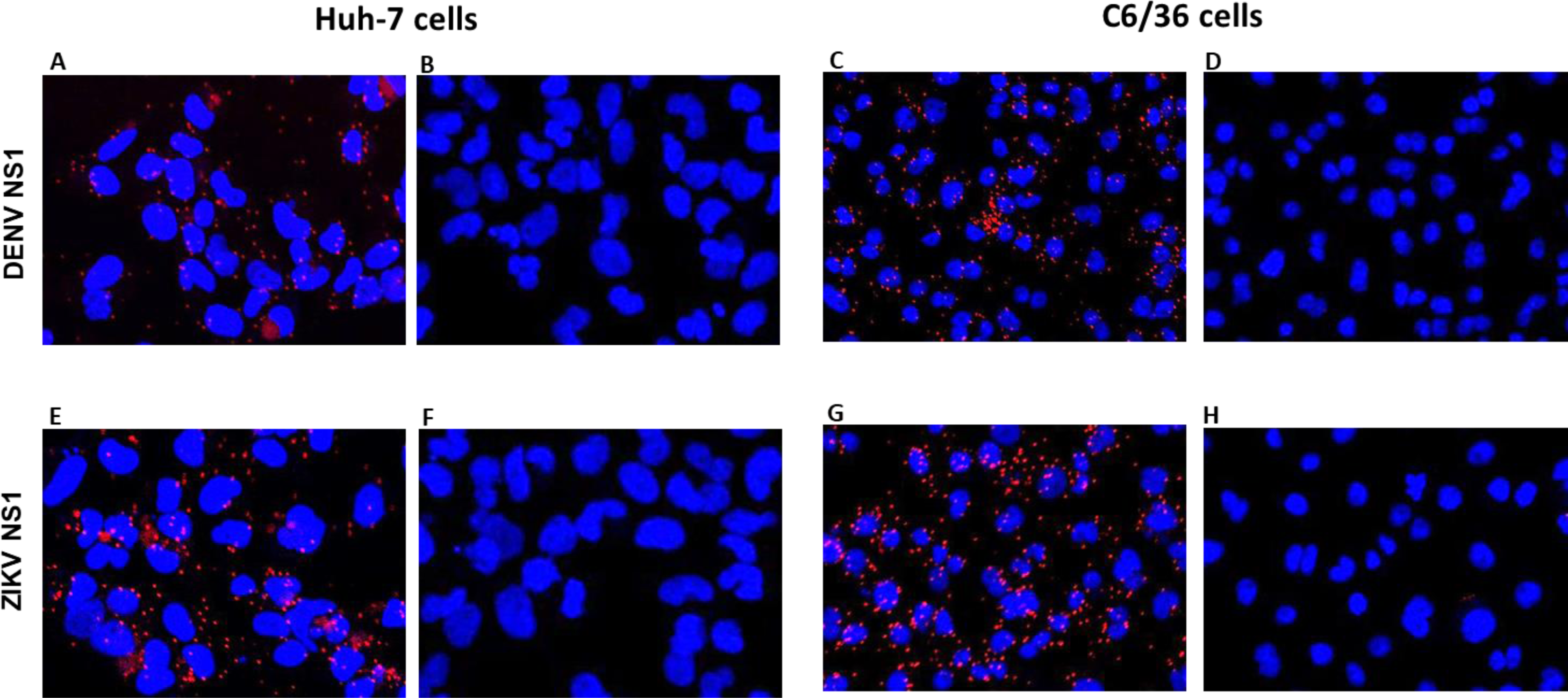
Proximity ligation assays for dengue and zika virus NS1 and the SRB1 in human and mosquito cells. Cells incubated with recombinant DENV NS1 (panels A, B, C and D) or ZIKV NS1 (panels E, F, G and H) at a concentration of 3,5 μg/well for 45 min, then fixed and processed following the manufacturer’s instructions. Positive interactions between NS1 and SRB1 are visualized as red dots. Cell nuclei were counterstained with DAPI. As negative controls, cells were preincubated with recombinant NS1, but primary anti-SRB1 antibody omitted (panels B, D, F and H). Experiments were carried out three times and typical results are shown.

The second approximation to evaluate DENV NS1 and SRB1 physical interactions was SPR. As shown in the sensorgram in Figure 8, the SRB1 developed a faster on-rate (0 to 200s) and slower off-rate (200s and on) with the interacting recombinant DENV NS1, at all the NS1 concentrations tested. The DENV NS1 showed a relative high affinity towards the SRB1, as indicated by the calculated average equilibrium dissociation constant (KD) values, ranging in the sub-nanomolar range (47.02 ± 0.01 nM versus 10mM or higher for non-specific binding interactions). Of note, the average equilibrium dissociation constant value observed for the recombinant NS1 and the SRB1 is in the same order of magnitude as those observed for the HDL and the SRB1 (KD=18.71 nM). The SPR results corroborate the results obtained in the infected cells by PLA, and together indicate that the SRB1 is bound by the DENV, and ZIKV NS1 protein.

**Figure 8.**
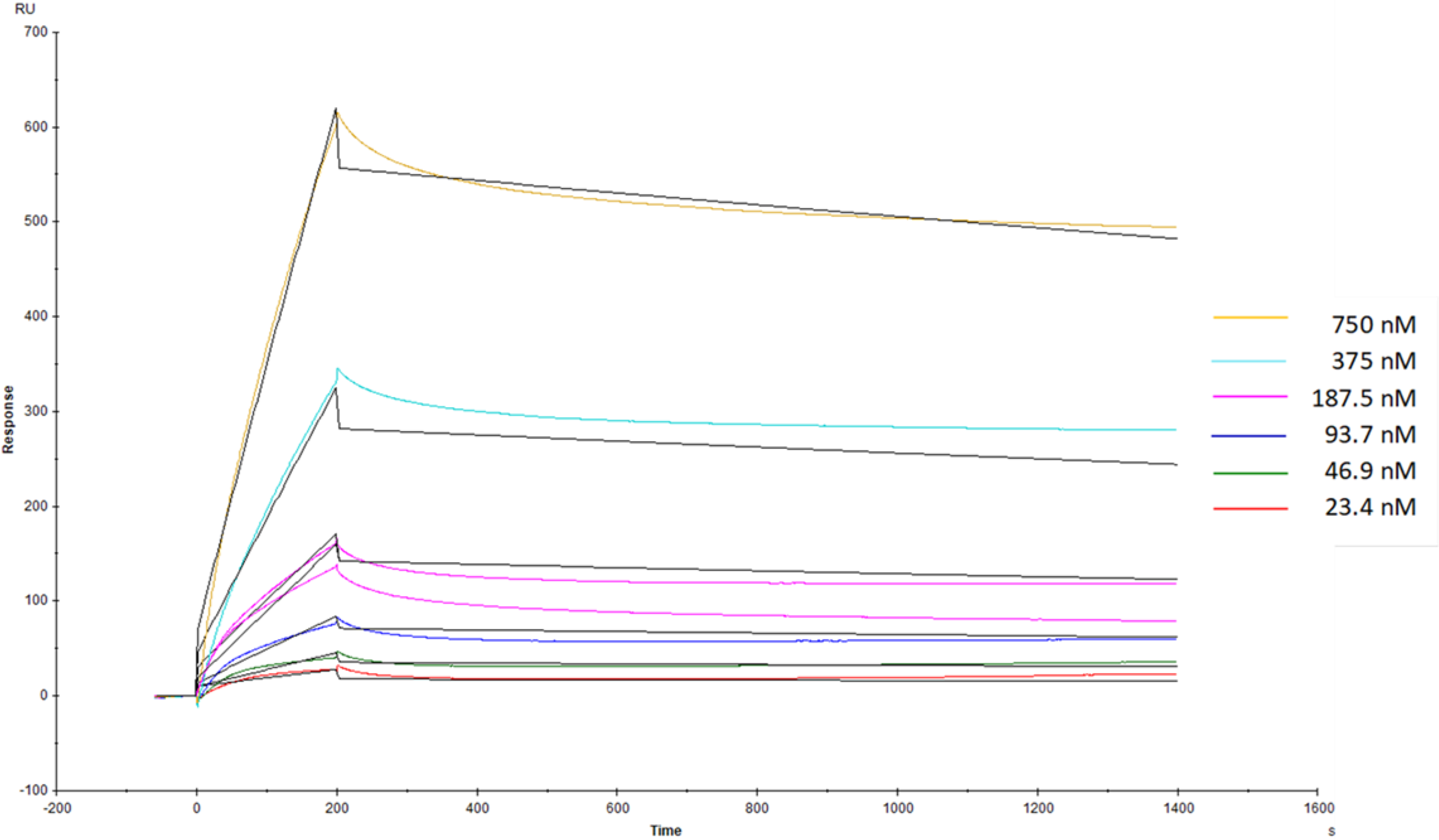
Surface plasmon resonance sensorgrams obtained for the binding interaction of NS1 with the SRB1. Increasing concentrations (color lines) of recombinant NS1 were tested for interaction with SRB1 immobilized on a Biacore sensor chip. Binding curves were expressed in resonance units (RU) as a function of time (seconds). Solid black lines through the curve show model fit for the calculation of interaction affinities.

## Discussion

NS1 is one of the most pleiotropic protein encoded by flaviviruses described so far. The ability of DENV NS1 to be secreted concomitantly with viral particles and reaching the extracellular milieu makes it a powerful “viral weapon” capable of participating in several pathogenesis processes, both intracellularly and extracellularly, thus favoring the installation and progression of the infection. DENV NS1 is internalized in cultured cells as well as *in vivo*, out of the context of the viral infection, to enhance viral production. Also, the endothelium disrupting capacity of flavivirus NS1 is dependent on cell internalization. However, little it is known about the mechanisms and host molecules involved in NS1 internalization. In this work, we assessed the participation of the SRB1 as a receptor of the DENV and ZIKV NS1 in human hepatic and insect cells. First, we observed the internalization of DENV and ZIKV NS1 in hepatic cells as previously reported (Alcon-Le Poder et al., 2005, Puerta-Guardo et al., 2019). Additionally, we observed that NS1 internalization also occurs in insect cells. Moreover, as has previously been reported for human hepatic and dendritic cells, preincubation of insect cells with DENV or ZIKV NS1 renders the cells more susceptible to the respective viral infection. The mechanisms by which NS1 contribute to increase the viral yield it is not known. However, it has been proposed that the accumulation of DENV NS1 in the late endosomal compartment after the entry to hepatocytes potentializes subsequent dengue virus infection *in vitro* (Alcon-Lepoder et al., 2005). In addition, the entry of NS1 to endothelial cells disrupt glycocalyx components which may favor virus binding and entry or potentially influencing virus dissemination and pathogenesis (Puerta-Guardo et al., 2016). In the mosquito cell, NS1 may also negatively regulate the innate cell response favoring viral replication (Liu et al., 2017). These mechanisms are not mutually excluding and may be all operating in the different cell types. In any case, additional experiments are needed to determine what are the mechanisms leading to increase DENV and ZIKV replication after NS1 cell intake.

The entry of a protein into a cell is a multifactorial process since the plasmatic membrane is a plethora of different kind of molecules determined by the origin of the cell, which in turn modulate the nature of proteins to be internalized (Walrant et al., 2017). In hepatic cells, SRB1 is the main receptor of the HDL, playing an important role in the homeostasis of lipoproteins circulating in serum (Shen et al 2019). Derived from the fact that NS1 is internalized in liver cells and is a lipoprotein in nature, in this work we present data indicating the participation of the SRB1 in the internalization process of DENV and ZIKV NS1 in hepatic and insect cells. Our results indicate that a functional SRB1-like receptor is encoded in the mosquito genome and expressed on the surface of the cell plasma membrane. As mentioned, the main function of the SRB1 is to participate in the influx and efflux of the HDL hauled molecules contributing to lipid homeostasis. Mosquito cells are unable to synthetize cholesterol *de novo* and SRB1-like receptor may participate in cholesterol uptake by these cells. On the other hand, our results showing the absence of the SRB1 on kidney derived Vero cells, agrees with previous results were the presence of NS1 was detected in the liver but not in the kidneys of mice exposed intravenously to DENV NS1 (Alcon-Lepoder et al.,2005).

The entry of DENV NS1 both in hepatic and insect cells was hampered by the preincubation with a specific monoclonal antibody against the SRB1. In both cells, there was a significant reduction of the NS1 internalization when 1 or 10 μg/mL of the antibody was used. However, the antibody only partially abolishes the NS1 binding and internalization, especially in hepatic cells, even at the higher concentrations used. It has been reported that position C323 plays a critical role in HDL binding and that the exoplasmic C384 plays a role in the selective HDL-cholesteryl esters (CE) uptake and in SRB1-mediated selective HDL-CE transport activity (Guo et al., 2011, Yu et al., 2011). The antibody used in this work was developed against an epitope near the amino-terminal region of the SRB1, away from positions C323 and C384. Thus, the antibody may only partially cause a steric impediment for NS1 internalization. Also, a significant but incomplete competition between NS1 and HDL, the natural ligand of the SRB, was observed. These results indicate that the SRB1 is used as a cell receptor by NS1. The participation of the SRB1 as cell binding molecule for NS1 was corroborated by the results obtained with transfected Vero cells where the sole expression of SRB1 is enough to render the cell susceptible to NS1 entry. Yet if HDL and NS1 bind to the same domains on the SRB1 molecule is unknown. In addition, as is the case for HCV entrance, where the successful binding in hepatic cells requires the SRB1, claudin-1, the human cluster differentiation CD81 and occludin as a minimal set of receptors working orchestrated as a multimolecular complex (Miao et al., 2017), SRB1 may not be the only receptor molecule for NS1.

To point a given molecule as a receptor for a ligand, evidence of direct interaction between the molecule and the ligand is required. In this work, direct physical interaction between the SRB1 and the DENV and ZIKV NS1 in Huh-7 and C6/36 cells was demonstrated by PLA. PLA is a relatively novel technique that uncover protein-protein interactions among proteins located not farther than 40 nm apart, and at variance with immunoprecipitation techniques, it does so maintaining the cell architecture (Sable et al., 2018). Moreover, PLA are run in parallel with several negative controls to assure the specificity of the reaction. Recently, the importance of NS1 present in the blood meal taken from the mammalian host to enhance the dissemination of ZIKV in the mosquito was reported (Liu et al., 2017).Thus, it is likely that the SRB1-like receptor expressed in mosquito cells is one of the molecules recognized by ZIKV NS1 upon blood ingestion. The PLA results for DENV NS1 and the SRB1 were corroborated *in vitro* by SPR. The results obtained by SPR clearly indicate physical interactions between the DENV NS1 and the SRB1 in a dose-dependent manner. Interestingly, the use of HDL in the SPR assays as a positive control showed that the affinity of the DENV NS1 and the human HDL for SRB1 does not differ significantly. This observation and the observation that HDL and DENV NS1 compete for the same receptor in Huh-7 cells, together with the high levels of circulating NS1 seen in patients during the acute phase of the disease, offers an explanation for the alterations in lipid homeostasis seen in dengue patients, as for example the decreased in HDL levels observed in patients with severe dengue (Li et al 2013).

In summary, in this work evidence is presented that the SRB1 in hepatic cells and an SRB1-like receptor in mosquito cells act as the cell binding protein for DENV and ZIKV NS1. Moreover, evidence is presented indicating that the C6/36 receptor is functional. It had been described that the selective uptake of cholesteryl esters by the cells through the SRB1, either in vitro and/or in vivo, do not show any specificity toward a single class of lipoproteins (Shen et al, 2018). Thus, we hypothesize that NS1 hijacks classic lipid routes for cell entry through the SRB1, but if the lipids carried out by NS1 are transferred to the cell is unknown. Once inside the cell, NS1 promotes viral replication and cause delocalization of tight junction proteins. Thus, future experiments should include the elucidation of the signaling pathways that are triggered after the SRB1 and NS1 interactions.

## Acknowledgments

This work was supported by UNAM.PAPIIT IT200418 to LAP and partially support by CONACYT (Mexico) grant 0254461 to JEL. Authors declare no conflict of interest.

